# Towards molecular-based functional classification of fetal bovine serum

**DOI:** 10.64898/2026.03.16.712020

**Authors:** Lara Magni, Natasja P. Christensen, Emmanuel Labaronne, Quan Shi, Lelde Berzina, Sandra Torres, Troels Kristiansen, Karsten Kristiansen

## Abstract

Quality and price of fetal bovine serum (FBS) are traditionally determined by geographical origin and parameters listed in the Certificate of Analysis (CoA). Despite its central role in cell culture, selecting suitable FBS batches remains costly and labor-intensive due to substantial batch-to-batch variation. We propose a molecular assessment strategy based on transcriptomic and cytokine profiling of cells cultured in different FBS batches to evaluate performance more reliably. Analysis of differential gene expression in three cell lines - MRC-5, Jurkat, and THP-1 - enables batch grouping and reveals pathway-specific effects, with immune-related pathways showing the most pronounced variability. Although CoA parameters can stratify batches by origin, they do not consistently correlate with cytokine secretion or gene expression across cell lines. These findings demonstrate that geographical origin is an inadequate predictor of functional FBS performance and that molecular profiling provides a more robust and informative assessment.

## Introduction

Fetal bovine serum (FBS) is a widely used supplement in cell culture, providing essential nutrients, growth factors, and hormones that support cell growth and survival. FBS is obtained from the blood of bovine fetuses after a series of processes including coagulation, centrifugation, and triple filtration, resulting in the sterile bioproduct used in cell culture ^1^. Despite its widespread use since the mid-20th century ^2^, the precise composition of FBS and its impact on cell growth and function remains poorly characterized, and batch variability continues to pose a significant challenge in scientific research ^1^. The batch-to-batch variations derive from several factors, such as the region of origin, production period, stage of fetal development, and production protocols ^3,4^. FBS remains a crucial cell supplement in research, diagnostics, pharmaceutical manufacturing, and numerous other biotechnological applications ^5-8^. However, the high degree of variability in FBS batches, even from the same manufacturer, remains a critical concern as it can compromise the reproducibility of experimental results within and between laboratories ^4,9,10^. In fact, a 2016 survey found that 70% of scientists encountered difficulties replicating experiments, often attributed to reagent variability, including FBS ^11^. Recent studies have highlighted how batch differences influence cell behavior, such as changes in ATP levels with consequent cell adaptation and variation in Ca^2+^-responses ^12^, IL-8 levels and inflammatory responses ^13^, redox biology ^14,15^. Nonetheless, comprehensive investigation of FBS batch variability remains limited. Currently, when FBS is recognized as a critical factor in an experimental setup, researchers often select the batch through a trial-and-error process that is both resource-intensive and costly ^4,6,16^. For a more general use, the selection process typically begins with choosing a batch based on its region of origin, which is also used as a criterion for assessing serum quality, with FBS from USA, Australia and New Zealand regarded as the highest quality.

In this study we performed an exploratory analysis on FBS batch variability. We initially validated a screening method based on transcriptome profiling using MRC-5 cells grown in four FBS batches. Then, we tested 20 distinct FBS batches of different geographical origin in three robust cell lines, MRC-5, Jurkat, and THP-1. We analyzed the transcriptome profile of cells grown with each FBS batch and we identified numerous differentially expressed genes (DEGs), even though these cell lines are considered quite robust exhibiting limited sensitivity to FBS batch variations. The results showed that transcriptome profiling enabled robust grouping of different batches of FBS which did not consistently correlate with geographical origin. We linked transcriptome profiles and secreted cytokine levels with biochemical parameters included in the Certificate of Analysis (CoA) provided by the manufacturer. Several parameters in the CoA clustered the FBS batches according to the area of origin. However, CoA parameters and consequently origin, did not robustly correlate with FBS performance as determined by transcriptome and cytokine profiles in the three selected cell lines. Our findings demonstrate that geographical origin is not a suitable predictor of functional FBS performance and that molecular profiling provides a more informative assessment.

In conclusion, we propose transcriptome profiles as an efficient methodology for FBS screening and potentially batch-matching. Such technology would increase the probability for customers of obtaining a new FBS batch with similar biological performance to the previous one. This would reduce the resources needed for batch testing and improving the reproducibility of research outcomes in biomedical and biotechnological fields.

## Methods

### Cell lines and media

The adherent human lung fibroblast cell line MRC-5 (CCL-171), the suspension human monocytic cell line THP-1 (TIB-202), and the human acute T-cell leukemic cell line Jurkat Clone E6-1 (TIB-152) were purchased from American Type Culture Collection (ATCC, USA). The mouse pre-adipocyte 3T3-L1 line was originally obtained from Professor M. Daniel Lane, Johns Hopkins University and propagated for years in our laboratory ^17^. The rat INS-1E cells were originally obtained from Professor Pierre Maechler, University Medical Center of Geneva ^18^. MRC-5 cells were grown in DMEM-GlutaMAX high glucose 25 mM (Thermo Fisher Scientific, 31966-047) supplemented with 10% fetal bovine serum (FBS) (Biowest, S00MZ2). THP-1 and Jurkat cells were maintained in RPMI (ATCC modified-Thermo Fisher Scientific, A1049101) supplemented with 10% FBS (Biowest, S00MZ2) under shaking conditions (Infors HT, Celltron, 110 rpm). The preadipocyte cell line 3T3-L1 was propagated in DMEM high glucose (Thermo Fisher Scientific, 52100021) supplemented with 10% calf serum (Fisher Scientific, B15-004) and 1% penicillin/streptomycin (p/s) (Gibco, 15140). The rat insulinoma INS-1E cell line was propagated in RPMI-GlutaMAX (Thermo Fisher Scientific, 61870-10) supplemented with 10% FBS, 50 μM 2-mercaptoethanol (Thermo Fisher Scientific, 31350-010), 1 mM sodium pyruvate (Sigma-Aldrich, S8636), 10mM HEPES (Thermo Fisher Scientific, 15630-106). All cell lines were kept in a humidified atmosphere at 37 °C with 5% CO2. Experiments were conducted using 20 different batches of FBS provided by Biowest (Table S1).

### 3T3-L1 differentiation, AdipoRed staining and quantification

Mouse pre-adipocytes 3T3-L1 cells were differentiated into mature adipocytes as follows: cells were seeded in 96-well plate in DMEM containing 10% calf serum. The medium was renewed every second day until the cells reached confluence. Differentiation was induced 2 days post-confluence (designated day 0) by changing the medium to DMEM supplemented with 10% FBS and 1 µg/ml insulin, 1µM dexamethasone, 0.5 mM methyl-isobutylxanthine (MIX). On day 2, the medium was changed to DMEM supplemented with 10% FBS and 1 µg/ml insulin. From day 4 to day 8, the medium was refreshed every second day with DMEM supplemented with 10% FBS. On day 8, the intracellular triglyceride content was detected using AdipoRed™ Assay Reagent (Lonza, PT-7009). The cells were washed with PBS and then incubated in 100 µl of a staining solution containing 2.5 µl AdipoRed (according to the manufacturer’s instruction) and 1µg/ml Hoechst 33342 (Life technologies, H3570) in PBS. Nuclear staining with Hoechst was used as a measure of cell number. Fluorescence for AdipoRed was measured using excitation at 485 nm with emission at 572 nm and for Hoechst using excitation at 350 nm with emission at 460 nm and quantified using a Varioskan LUX Multimode Microplate Reader (Thermo Fisher Scientific). Images of stained cells were obtained using an Opera Phenix™ Plus high-content imaging system (Revvity). AdipoRed intensity was normalized to cell number. A FBS batch previously validated for adipocyte differentiation was used as control. The samples were considered normally distributed, as they passed the Shapiro–Wilk test, and the statistical analysis was performed with one-sample t test.

### Insulin secretion from INS-1E cells

Rat insulinoma INS-1E cells were seeded into a 48-well plate at a final density of 50.000 cells/well and allowed to adhere and grow undisturbed for 72 h. Prior to the experiment, the cells were kept for 2 h in Krebs-Ringer buffer (KRB) with the following composition: 135 mM NaCl, 3.6 mM KCl, 5 mM NAHCO_3,_ 0.5 mM NaH_2_PO_4_.H_2_O, 0.5 mM MgCl_2_.6H_2_O, 1.5 mM CaCl_2_.2H_2_O, and 10 mM HEPES, pH 7.4, BSA (0.2%), and 5.5 mM glucose. The cells were stimulated for 30 min with either 2.8 mM or 16.7 mM glucose in KRB and the cell supernatant was collected stored at −80 °C until measurement of insulin. Samples were analyzed using MSD Mouse/Rat Insulin Kit (Mesoscale, K152BZC) according to the manufacturer’s instructions. The values of secreted insulin at low (2.8 mM) glucose and high (16.7 mM) glucose were normalized to the average of the secreted amount at 2.8 mM glucose of each FBS. The samples were considered normally distributed, as they passed the Shapiro–Wilk test, and therefore, the two secretions were compared for each FBS batch with t-test analysis.

### Samples collection and RNA extraction

MRC-5, THP-1 and Jurkat cells were seeded in triplicates in 6-well plate and grown in each of the 20 different FBS batches. The MRC-5 cells were grown for 6 days until confluence, Jurkat and THP-1 were allowed to grow for three and four days, respectively, and harvested in the exponential growth phase. The supernatant was collected and stored at -80°C and the cells were harvested in TRIzol reagent and stored at -20 °C until extraction. RNA extraction was performed using the Direct-zol RNA Miniprep kit (Zymo Research, R2053), and extracted RNA was eluted in 30 µl DNase/RNase-free distilled water (Invitrogen, 10977035) and stored at -80 °C until further processing.

### Library preparation and RNA sequencing

The quantity and quality of the extracted RNA were determined using the Qubit RNA BR Assay Kit (Thermo Fisher Scientific, Q10211) and the High Sensitivity RNA Assay Kit (Agilent Technologies, 5067-5579), respectively. The integrity of RNA was evaluated by RNA Integrity number (RINe) on a 4200 TapeStation system (Agilent Technologies). For messenger RNA (mRNA) enrichment, 300 ng of total RNA were processed using the Dynabeads mRNA Purification Kit (Thermo Fisher Scientific, 61006). RNA library preparation was made using the MGIEasy RNA Directional Library Prep Set (MGI Tech Co., Ltd, 1000006386) following the manufacturer’s instructions. Concentration and fragment analysis of prepared libraries were performed using the Qubit 1X dsDNA HS Assay Kit (Thermo Fisher Scientific, Q33231) and Agilent High Sensitivity D1000 Assay Kit (Agilent Technologies, 5067-5584 and 5067-5585). Pooled and concentrated libraries were sequenced on a DNBSEQ-G400 sequencer (MGI Tech Co., Ltd.) using the DNBSEQ-G400RS High-throughput Sequencing Set (FCL PE100) (MGI Tech Co., Ltd., 1000016952) according to the manufacturer’s instructions.

### Read processing

RNA sequencing data processing and analysis were performed using a custom Nextflow ^19^ pipeline. Briefly, raw sequencing data quality was initially assessed using FastQC v0.12.1. Read preprocessing was performed using fastp v0.20.1 ^20^ with a qualified Phred quality score threshold of 20. The preprocessing parameters included poly-X trimming, automatic adapter detection for paired-end reads, and disabled length filtering. Processed reads were filtered against rRNA and snRNA sequences using Bowtie2 v2.3.4.1 ^21^ with the ‘--very-sensitive’ preset to minimize contamination. Unaligned reads were retained for subsequent analysis. The filtered reads were aligned to the GRCh38.p12 human reference genome using STAR v2.7.3a ^22^. STAR indices were generated using the reference genome and GENCODE v30 annotation file. The mapping was performed with the following parameters: --outFilterMultimapNmax 20 --alignSJoverhangMin 8 --alignSJDBoverhangMin 1 -- outFilterMismatchNmax 999 --outFilterMismatchNoverReadLmax 0.04 --alignIntronMin 20 --alignIntronMax 1000000 --alignMatesGapMax 1000000 --quantMode TranscriptomeSAM, with quantification of both genome-aligned reads and transcriptome-aligned reads. For transcript quantification, Salmon v1.10.3 ^23^ was employed in alignment-based mode using the mapped transcriptome alignments. The quantification was performed against the GENCODE v30 transcriptome reference with --libType A --gencode parameters. A transcript-to-gene mapping file was generated from the GENCODE v30 GTF annotation to facilitate gene-level abundance estimates in downstream analyses. All processing steps were executed in a containerized environment using Docker to maintain software version consistency.

### Differential gene expression analysis

Gene expression analysis was performed in R (v4.3.3) using the tximport v1.30.0 package to import Salmon quantifications and DESeq2 v1.42.1 ^24^. Prior to differential expression analysis, lowly-expressed genes were filtered using a custom filtering approach that retained genes with a minimum CPM of 1 in at least 3 samples. Count data normalization and differential expression analysis were performed using DESeq2’s model ∼ FBS. Genes with an adjusted p-value < 0.05 and absolute log2 fold change > log2(1.5) were considered significantly differentially expressed.

### sPLS and PCA

For RNAseq data, sparse Partial Least Squares (sPLS) analysis was performed on variance-stabilized transformed (VST) count data to assess sample relationships and identify potential batch effects. Additionally, variance-based filtering was applied to retain genes with variances above 1e-5 after VST. SPLS as performed using mixOmics v6.26.0 ^25^, package. For analysis of FBS biochemical parameters, Principal Component Analysis (PCA) was performed using the PCAtools v2.14.0 package. Data was scaled and centered prior to analysis to account for differences in measurement units across components. AREA annotation was added manually.

### Network analysis of differentially expressed genes

The relationships between different FBS batches were visualized through a network approach. Briefly, from the differential expression analysis, the number of differentially expressed genes (DEGs) (adjusted p-value < 0.05, |log2FC| > log2(1.5)) was used to calculate a similarity score between conditions using the formula: similarity = 1/(1 + total_DEGs). The network was constructed using batches as nodes and edges representing similarity between batches (weighted by similarity score). Community detection was performed using the Louvian algorithm with a resolution of 0.75. Visualization was done using tidygraph and ggraph packages.

### Pathways analysis

Gene set enrichment analysis was performed to identify biological pathways affected by different FBS batches. The Molecular Signatures Database (MSigDB) Hallmark gene set collection was used to identify enriched pathways. Enrichment analysis was performed using the clusterProfiler v4.10.1 package ^26^ with over-representation analysis (ORA) method and using Benjamini-Hochberg correction for multiple testing. A network-based approach was used to identify relationships between enriched pathways. For each significant Hallmark pathway across all comparisons, Jaccard similarity was calculated between pathway pairs based on co-occurrence. Edges were created between pathways with similarity above a threshold (0.3). The Community analysis was performed using the Louvain algorithm with a resolution of 0.7. The network was visualized using a force-directed layout where nodes represent Hallmark pathways; nodes size indicates the frequency of pathway occurrence across comparisons; node color represents community membership; edge width represents similarity between pathways.

### FBS Cluster Assignment

FBS clusters were defined based on hierarchical clustering of the number of differentially expressed genes (defined as absolute log2 fold Change > 0.6 and adjusted p-value < 0.05) using a threshold of 100 differentially expressed genes (DEGs). Samples were assigned to clusters as follows: MRC-5 cells: Cluster 1 (FBS.13), Cluster 2 (FBS.01, 02, 03, 04, 05, 06, 07, 08, 09, 10, 20), Cluster 3 (FBS.14, 15, 16), Cluster 4 (FBS.17). Jurkat cells: Cluster 1 (remaining FBS), Cluster 2 (FBS.11, 15, 16), Cluster 3 (FBS.05, 13), Cluster 4 (FBS.03, 06, 08, 17, 18, 19). THP-1 cells: Cluster 1 (FBS.14, 15, 16), Cluster 2 (FBS.07, 11, 13), Cluster 3 (FBS.03, 04, 12), Cluster 4 (remaining FBS).

### Weighted Gene Co-expression Network Analysis

Weighted Gene Co-expression Network Analysis (WGCNA) was performed using the WGCNA package (v1.72-5) in R to identify co-expressed gene modules and relate them to FBS cluster phenotypes. Analysis was conducted separately for each cell type using DESeq2 results. Expression data were prepared using VST from DESeq2 objects. Soft-thresholding power was determined using pickSoftThreshold (range 1-20), selecting the lowest power achieving scale-free topology R^2^ ≥ 0.8. Networks were constructed using blockwiseModules with parameters: networkType=“signed”, correlationType=“pearson”, minModuleSize=30, mergeCutHeight=0.25. Module eigengenes were calculated using moduleEigengenes, and Pearson correlations with FBS clusters were computed. Significant relationships were defined as p < 0.05 using corPvalueStudent. GO enrichment analysis was performed using clusterProfiler (v4.10.1) with enrichGO function (pvalueCutoff=0.05, qvalueCutoff=0.2, Benjamini-Hochberg correction).

### Cytokine release

Cells supernatants were analyzed to quantify 20 different cytokines/factors. The samples were obtained from cells grown with the 20 different FBS batches in triplicates and assayed with the MSD V-PLEX Cytokine Panel 1 Human Kit (Mesoscale K15050D) and the MSD V-PLEX Proinflammatory Panel 1 Human Kit (Mesoscale K15049D). Control samples composed of medium and 10% FBS (of each batch) were also included in the assay. Only the cytokines/factors detectable in all samples were included in the further analysis. Cytokine abundances were visualized as mean +/- SEM of biological replicates. Statistical differences were tested using Wilcoxon Rank-Sum test, followed by a Benjamini-Hochberg (BH) correction for multiple testing.

### Correlation analysis

For each cell line, Spearman correlation coefficients were calculated between cytokine levels and Certificate of Analysis (CoA) parameters using the cor function in R. Correlation matrices were visualized using the pheatmap package. For the correlation between the transcriptome and the CoA parameters, two enrichment methods were applied to analyze pathway activity, GSVA (Gene Set Variation Analysis) and ssGSEA (single-sample Gene Set Enrichment Analysis), both implemented via the GSVA package with Gaussian kernel distribution. Pathway activity scores were correlated with FBS biochemical parameters using Pearson correlation. The resulting correlation matrices were visualized using the pheatmap package. The complete R analysis code and associated functions are available at https://gitlab.com/adlin-science-public/fbs-molecular-classification.git

### Statistics and reproducibility

All statistical analyses were performed using R version 4.3.3 or Prism9 software (GraphPad). RNA sequencing data processing utilized a custom Nextflow pipeline with FastQC v0.12.1, fastp v0.20.1, Bowtie2 v2.3.4.1, STAR v2.7.3a, and Salmon v1.10.3. Statistical tests and statistical significance are described in the in the corresponding paragraph and/ or in the figure legend. For functional assays, normality was assessed using Shapiro-Wilk test, and statistical comparisons were performed using one-sample t-test (3T3-L1 differentiation), paired t-test (insulin secretion), or Wilcoxon Rank-Sum test with Benjamini-Hochberg correction (cytokine analysis). RNA samples and functional experiments were conducted in three independent wells. The validation of a FBS screening workflow for reproducibility was assessed by analyzing six independent experiments, including samples from three independent wells.

## Results

### Differential effects of FBS batches on adipocyte differentiation and insulin secretion

We selected 3T3-L1 cells and INS-1E cells as examples of sensitive models to test the effect of different FBS batches in adipocyte differentiation and insulin secretion, respectively. We tested 16 FBS batches for differentiation of 3T3-L1 cells, a cell line known to be sensitive to FBS variation ^27^, to evaluate variation of differentiation between the selected FBS batches. As expected we observed significant differences in the ability of the different FBS batches to support adipocyte differentiation using a commonly used protocol including treatment with dexamethasone, 3-isobutyl-1-methylxanthine, and insulin ^17^. The level of differentiation was assessed qualitatively via live cell imaging (Figure 1A). In addition, the degree of differentiation was evaluated measuring triglyceride accumulation with AdipoRed fluorescence normalized to cell number (Hoechst staining) (Figure 1B). A clear variation in differentiation was observed, ranging from 115% to 66% relative to a previously selected control FBS batch (denoted as CTR) based on AdipoRed fluorescence (Figure 1B). Moreover, we tested 5 different FBS batches in INS-1E cells (Figure 1C). The cells were grown for up to 6 weeks in media containing 10% of each FBS batch and the insulin secretion capability was assessed at week 2 and 6. The secreted insulin showed an interesting trend between week 2 and 6. At week 2 all the FBS batches tested supported induction of insulin secretion in response to a high physiological level of 16.7 mM glucose. However, at week 6, FBS 1, 11, 13, and 15 exhibited a loss of glucose-induced insulin secretion, while FBS 12 maintained and even increased glucose-induced insulin secretion from around 3 folds in week 2 to 15 folds in week 6. In conclusion, the tested FBS batches showed very different capabilities of promoting adipocytes differentiation and insulin secretion providing a clear example of the importance of FBS batch selection for cell-based experiments. Based on these findings, we conducted a more in-depth analysis of the FBS batches to further investigate to what extent different FBS batches impacted transcriptional profiles and cytokine expression.

**Figure 1.**
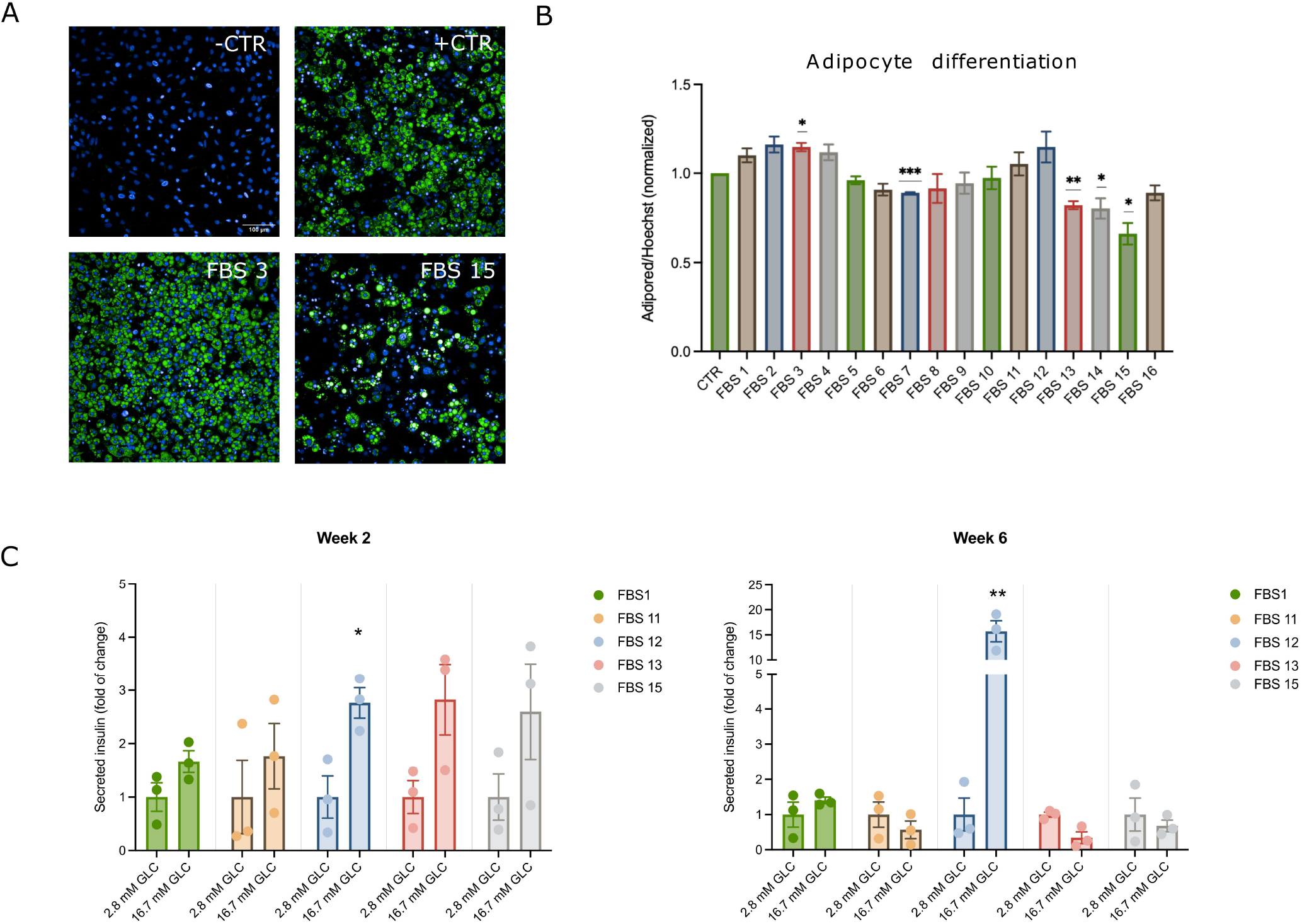
FBS batch affects adipocyte differentiation and insulin secretion. (A) Live cell imaging of differentiated 3T3-L1 cells. AdipoRed staining was used to qualitatively assess adipocyte differentiation. Cell nuclei were stained with Hoechst. Pictures of cells differentiated with 4 FBS batches have been chosen as representative of the FBS effect on adipocytes differentiation. The scale bar corresponds to 100 μm. (B) Quantification of 3T3-L1 differentiation. AdipoRed fluorescence was quantified and normalized first on cell number (Hoechst fluorescence). The differentiation is normalized to the one obtained with to a previously qualified FBS batch denoted as CTR. Statistical analysis was performed with one sample t-test and significance is indicated as *p < 0.05, **p < 0.01, ***p < 0.001. (C) Insulin secretion of INS-1E cells cultured for 2 weeks and 6 weeks in different FBS batches. Insulin secretion after stimulation with 16.7 mM glucose is reported as fold of change compared to the amount secreted with 2.8 mM glucose. The values of secreted insulin at 2.8 mM glucose and 16.7 mM glucose were normalized to the average of the secreted amount at 2.8 mM glucose of each FBS. The statistical analysis was performed with t-test and compared the low and high glucose secretion of each FBS. Significance is indicated as *p < 0.05, **p < 0.01. The experiment was performed once with three technical replicates.

### Validation of a FBS screening workflow for reproducibility

To delve deeper into the sources of variability between different serum batches we examined to what extent differences in transcriptome patterns could be used as a sensitive fingerprint to characterize differences between serum batches. Initially, we screened 8 cell lines and selected the stable and widely used MRC-5 cells to evaluate whether the results obtained with a given FBS batch were reproducible. Cells were seeded and allowed to grow until one day post-confluence using 4 different FBS batches. Each serum batch was tested in six independent biological replicates, each analyzed in triplicate. RNA was extracted and subjected to RNA sequencing providing at least 10 million reads per sample. The data were analyzed using sparse partial least squares (sPLS) regression revealing batch-related differences, with biological replicates clustering consistently within their respective batch groups (Figure S1A). Moreover, the correlation coefficients of the samples belonging to each batch group were high and similar across the biological replicates (Figure S1B). We conclude that the experimental variation between replicates was minimal, and FBS batch identity contributed significantly to transcriptomic variance, rather than procedural or technical errors.

### The number and types of Differentially Expressed Genes (DEGs) reflect batch differences

To evaluate more closely differences introduced by FBS batch variations, we selected three stable and robust cell lines, MRC-5, Jurkat, and THP-1 and analyzed the transcriptome profiles after exposure to each of 20 different FBS batches. The 20 FBS batches covered different regions of origin (Table S1). Each batch was tested in three technical replicates for each cell line. The subsequent analysis of differentially expressed genes (DEGs) revealed clear differences between the individual serum batches. Each gene was classified as a DEG when the difference in the expression had a p value <0.05 and exhibited a minimum 1.5-fold of change (Table S2). To obtain a global view of the similarity between FBS batches, we performed network analysis using the inverse of the number of DEGs as a similarity metric (Figure 2A). Consequently, two FBS batches positioned close together in the network represent FBS that differed by few DEGs.

**Figure 2.**
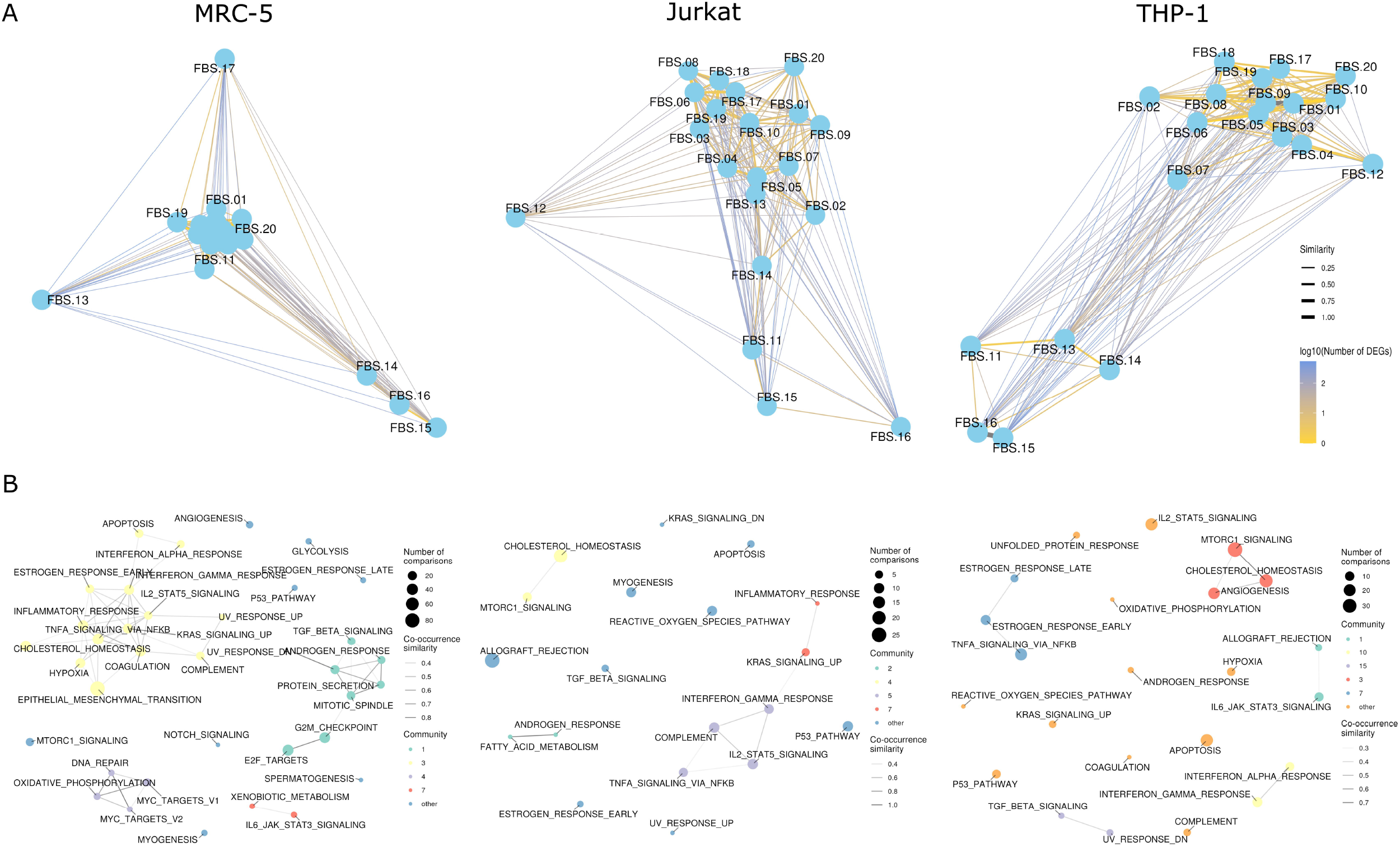
FBS batch comparison reveal cell type-specific transcriptional networks. (A) Differential expression similarity networks. Network representations for each cell type showing pairwise relationships between FBS batches based on differential gene expression patterns. Nodes represent FBS batches, edge width indicates similarity (inverse of number of DEG count) and edge color shows log10 (number of DEGs). Networks were constructed using stress layout. Nodes close together indicates similar FBS batches (few DEGs between conditions). (B) Hallmark pathway co-enrichment networks. Networks displaying relationships between significantly enriched Hallmark pathways across FBS comparison for each cell type. Nodes represent enriched pathways appearing in at least two comparisons, node sizes indicate frequency across comparisons, and edges connect pathways with Jaccard similarity > 3. Communities detected using Louvain clustering. Single-node communities are grouped as “other”. Highlighted with a blue square is the inflammatory response pathway.

Most of the serum batches seemed to cluster independently of the cell lines, a small number of serum batches exhibited distinct properties, in particular FBS 15 and 16, and few serum batches showed a cell type specificity in clustering. To delve deeper into the differences between the serum batches, we used Gene Set Enrichment Analysis (GSEA) of the DEGs based on hallmark gene sets from MSigDB (v7.5.1) to explore which pathways were most affected by different FBS batches (Figure 2B). The analysis showed that MRC-5 exhibited the highest number of differentially affected pathways in response to different serum batches, followed in order by THP-1 and Jurkat. In fact, in MRC-5 cells 80 pathways were differentially enriched depending on the different serum batches, including pathways involved in epithelial to mesenchymal transition. In THP-1 cells 30 pathways were differentially enriched, whereas for Jurkat cells - 25 pathways. Overall, in all three cell lines, pathway related to inflammation and immune responses seemed to be the most affected by different FBS batches.

### WGCNA analysis reveals cell type-specific gene expression modules associated with FBS-dependent clusters

To further characterize molecular signatures underlying the FBS batch groupings identified through differential expression analysis, we performed Weighted Gene Co-expression Network Analysis (WGCNA). Based on hierarchical clustering using the DEGs as a distance metric, we established distinct FBS clusters using a threshold of 100 DEGs, below which conditions were considered similar (Figure 3A). This approach identified 4 distinct FBS clusters in both Jurkat and THP-1 cells, and 5 clusters in MRC5 cells, revealing cell type-specific responses to different FBS batches.

**Figure 3.**
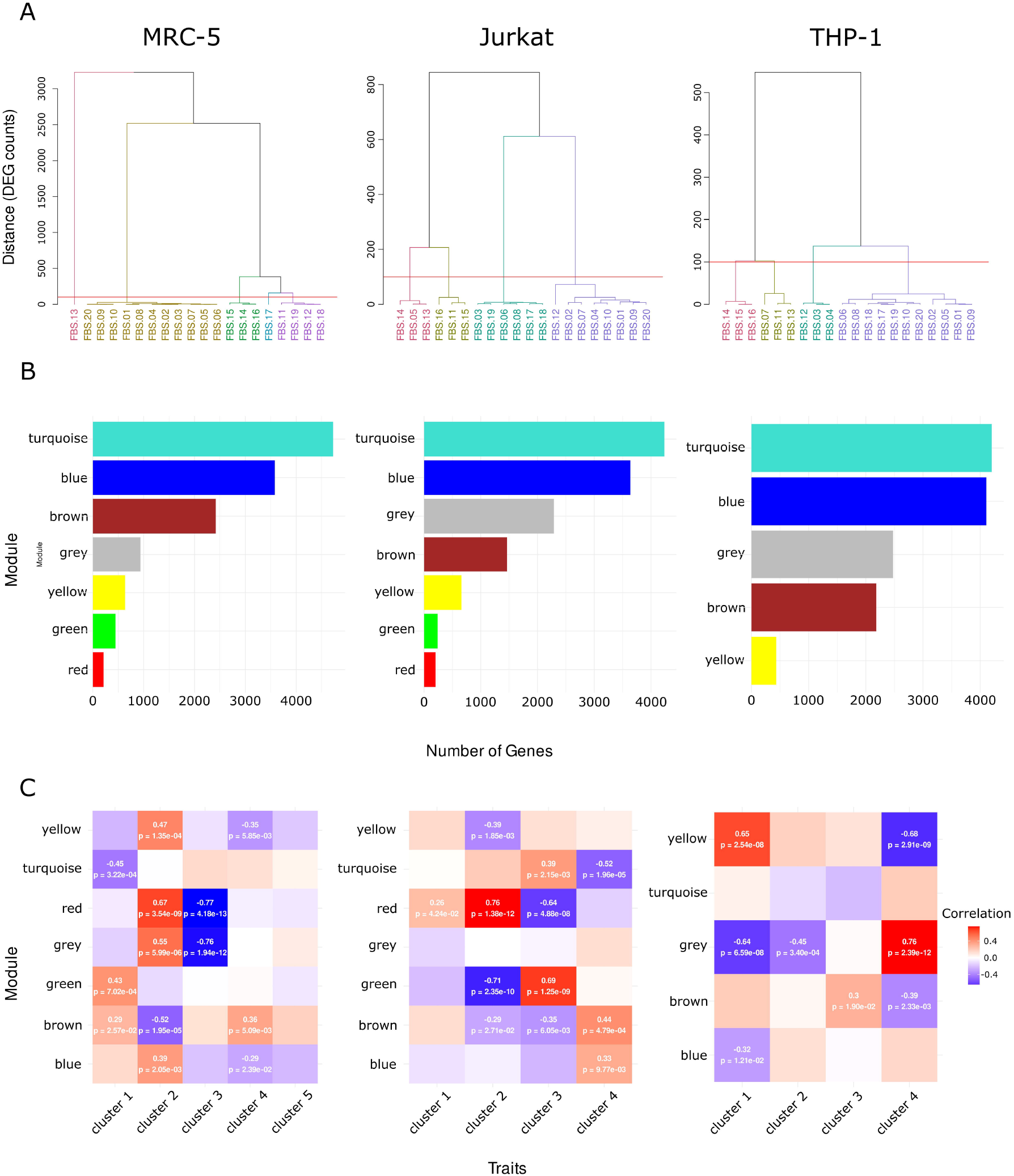
WGCNA analysis reveals cell type-specific gene expression modules associated with FBS-dependent clusters. (A) Hierarchical clustering dendrograms of FBS conditions based on differential gene expression profiles in MRC-5 (left), Jurkat (middle), and THP-1 (right) cells. Clustering was performed using the number of differentially expressed genes (DEGs; |log2FC| > 0.6, adjusted p-value < 0.05) as a distance metric. Horizontal red lines indicate the clustering threshold of 100 DEGs, defining distinct FBS clusters. MRC-5 and THP-1 cells show 4 distinct clusters, while Jurkat cells display 5 clusters, revealing cell type-specific responses to FBS alternatives. (B) Module size distributions from WGCNA for MRC-5 (left), Jurkat (middle), and THP-1 (right) cells. Bar colors represent distinct co-expression modules identified using signed networks with Pearson correlation (soft-thresholding power selected to achieve scale-free topology R^2^ ≥ 0.8; minimum module size = 30 genes; merge cut height = 0.25). MRC-5 cells: 7 modules (turquoise: 4,728 genes; blue: 3,580; brown: 2,417; grey: 936; yellow: 634; green: 447; red: 234). Jurkat cells: 7 modules (turquoise: 4,234; blue: 3,635; grey: 2,289; brown: 1,464; yellow: 660; green: 240; red: 204). THP-1 cells: 5 modules (turquoise: 4,201; blue: 4,16; grey: 2,478; brown: 2,184; yellow: 434). (C) Module-trait relationship heatmaps showing Pearson correlations between module eigengenes and FBS cluster assignments for MRC-5 (left), Jurkat (middle), and THP-1 (right) cells. Each cell displays the correlation coefficient (color scale: blue = -0.5, white = 0, red = 0.5) and corresponding p-value. Significant correlations (p < 0.05, calculated using corPvalueStudent) indicate modules with expression patterns specifically associated with FBS clusters. Notable associations include: MRC-5 turquoise module with Cluster 1 (r = - 0.45, p = 3.22×10 ); Jurkat red module with Cluster 2 (r = 0.76, p = 1.38×10 ^12^); THP-1 yellow module with Cluster 1 (r = 0.65, p = 2.54×10 8).

The WGCNA analysis successfully identified co-expression modules that captured the molecular basis of these cluster distinctions (Figure 3B). In MRC-5 cells, 7 gene modules were detected (turquoise: 4,728 genes; blue: 3,580 genes; brown: 2,417 genes; grey: 936 genes; yellow: 634 genes; green: 447 genes; red: 212 genes), while Jurkat cells showed 7 distinct modules (turquoise: 4,234 genes; blue: 3,635 genes; grey : 2,289 genes; brown: 1,464 genes; yellow: 660 genes; green : 240 genes ; red: 204 genes), and THP-1 cells exhibited 5 modules (turquoise: 4,201 genes; blue: 4,016 genes; grey: 2,478 genes; brown: 2,184 genes; yellow: 434 genes). These modules represent groups of genes with similar expression patterns across samples, potentially reflecting coordinated biological processes.

A cell type-specific module-cluster associations revealed distinct molecular programs with FBS batch-specific effects (Figure 3C). In MRC-5 cells, several modules exhibited strong associations with specific FBS clusters. Cluster 1, defined solely by FBS.13, was negatively correlated with the turquoise module (r = -0.45, p = 3.22×10 □ □ ), enriched for cell cycle regulation, chromosome segregation, and DNA replication processes, suggesting marked inhibition of proliferative programs unique to this batch. Cluster 4, corresponding to FBS.17 alone, displayed distinct correlations with the brown module (r = 0.36, p = 5.09×10□^3^) related to mitochondrial function and ribosome biogenesis. By contrast, the majority of FBS (cluster 2, comprising 11 batches: FBS.09, 04, 02, 03, 07, 05, 06, 01, 08, 10, 20) induced coherent signatures across modules linked to mesenchymal differentiation, immune receptor recombination, and chromatin organization, suggesting a relatively homogeneous response. Interestingly, a subgroup of FBS (14, 15, 16 forming cluster 3) consistently separated from the bulk, showing strong negative correlations with both red (r = -0.77, p = 4.18×10 □^13^) and grey (r = -0.76, p = 1.94×10 □^12^) modules, suggesting reduced mesenchymal and chromatin organization signatures.

In Jurkat cells, FBS-induced clustering was primarily driven by immune-related pathways. Cluster 2 (FBS.16, 15, 11) correlated strongly with the red module (r = 0.76, p = 1.36×10 □^12^), enriched for B cell activation, glial cell differentiation, and MHC class Ib antigen presentation, while showing strong negative correlation with the green module (r = -0.71, p = 2.35×10 □^1^□), involved in T cell-mediated immunity and adaptive immune responses. Conversely, cluster 3 (FBS.06, 08, 03, 19, 17, 18) displayed the opposite trend, with strong positive association with the green module (r = 0.69, p = 1.25×10□ □) and negative correlation with the red module (r = -0.64, p = 4.88×10 ), highlighting FBS-dependent shifts between immunomodulatory and cytotoxic programs. Additionally, cluster 4 (remaining FBS batches) was positively correlated with the brown module (r = 0.44, p = 4.79×10□4) involved in mitochondrial function and negatively with the turquoise module (r = -0.52, p = 1.96×10□5) related to growth regulation and autophagy.

In THP-1 cells, FBS variation predominantly impacted metabolic versus differentiation signatures. Cluster 1 (FBS.14, 15, 16) was positively associated with the yellow module (r = 0.65, p = 2.54×10□8), enriched for small molecule catabolism, glycolysis, and cholesterol biosynthesis, and strongly negatively correlated with the grey module (r = -0.64, p = 6.59×10 □8), involved in hematopoiesis regulation and myeloid cell differentiation. In contrast, cluster 4 (majority of remaining FBS) showed the inverse pattern, with strong positive correlation with the grey module (r = 0.76, p = 2.39×10 □^1^2) and negative correlation with the yellow module (r = -0.68, p = 2.91×10 □9), consistent with a differentiation-oriented profile. Cluster 2 (FBS.07, 11, 13) and cluster 3 (FBS.12, 03, 04) showed intermediate associations, with cluster 3 positively correlated with the brown module (r = 0.3, p = 1.90x10^-2^) related to mitochondrial function.

### Cross-cell line analysis reveals batch-specific outliers and conserved FBS subgroups

Notably, FBS.13 and FBS.17 formed unique single-lot clusters exclusively in MRC-5 cells, displaying distinct transcriptional signatures not observed in Jurkat or THP-1 cell lines, suggesting cell type-restricted effects of these batches. Conversely, a consistent subgroup of FBS (14, 15, 16) segregated together across multiple cell types: forming cluster 3 in MRC-5 (associated with reduced mesenchymal signatures), partially represented in Jurkat cluster 2 (FBS.15, 16 with immunomodulatory profiles), and constituting cluster 1 in THP-1 (associated with enhanced metabolic activity) (Figure 3). This cross-lineage consistency suggests shared biological properties of these FBS batches that transcend cell type-specific responses.

### Serum batches differentially affect levels of secreted cytokine

Since inflammation and immune-response pathways were highlighted in all three cell lines by the DEGs analysis (Fig 2B), we selected a panel of 20 cytokines that were measured in the supernatant of the same cells used for the transcriptome analysis. Although all cytokines were detected in at least one FBS batch for each cell line, we decided to focus on cytokines that were detectable across all FBS batches. A total of 13 cytokines were detected in the supernatant of MRC-5, while three cytokines were detectable for Jurkat and THP-1 cells (Figure S2). Interestingly, the different cytokines were affected by FBS batches to different extents. Some cytokines, such as IL-8, IL-6, IL-15 and GM-CSF in MRC-5, IL-16 and TNF-β in Jurkat and VEGF in THP-1 cells, were present in the cell supernatant in relatively high concentrations and/or strongly affected by different FBS batches, while others with levels slightly above the detection limits were less sensitive to FBS batch variation. However, comparing cytokines detected in more than one cell line, such as IL-2 for Jurkat and MRC-5, IL-8 for THP-1 and MRC-5, and IL-16 for Jurkat and THP-1 we noticed that, although affected by FBS batch, the extent of the induction/inhibition was not consistent within the same FBS batch. Nonetheless, cytokine levels clearly depended on the FBS batch, but not consistently across the cell lines or across batches with the same origin.

### Certificate of Analysis (CoA)-based clustering of FBS batches according to origin and impact on molecular performance

The CoA is a document provided for each FBS batch that includes biological and biochemical parameters measured in the serum. We selected all the quantifiable and absolute measurements of CoAs for a total of 31 parameters and performed a PCA analysis. The resulting PCA plot illustrated clear clustering of the batches based on the geographical region of origin. Specifically, the data showed that FBS from distinct regions formed four separate clusters, consisting of Latin America, Europe plus South Africa, Ireland, and Chile (Figure 4A). The variation in the levels of selenium, chloride, iron, glucose, uric acid, phosphorus, and potassium contributed to the separation (Figure 4B). The first two principal components, which together explained around 50% of the total variance, primarily captured the differences in FBS composition across regions. The clustering pattern observed in the plot suggests that the identified biochemical parameters serve as relevant distinguishing factors between the regions from which the FBS originated. This clustering highlights the potential influence of regional agricultural practices and environmental factors. However, it is not known if CoA parameters impact FBS performance, which would justify an origin-based FBS classification. Therefore, we performed a correlation of the CoAs first with the detectable cytokine levels (Figure S3A) and next with the transcriptome profiles (Figure S3B). The correlation between CoAs and cytokines showed that the same cytokines did not correlate consistently with CoA parameters in the different cell lines. However, we observed significant correlations in each cell line with MRC-5 cells showing a strong correlation between pro-inflammatory cytokines and CoA parameters. Here, an interesting tendency was observed for IL-8, IL-1β and TNFα, which exhibited positive correlations with parameters such as glucose, iron, uric acid, and phosphorus that drive the FBS batch clustering towards a European origin, while exhibiting inverse correlation to selenium, which drives the clustering towards Latin America. In Jurkat and THP-1 cells, only few significant correlations between cytokines and CoA parameters were observed and these were in general not linked with parameters that drive the FBS batch origin clustering (Figure S3A).

**Figure 4.**
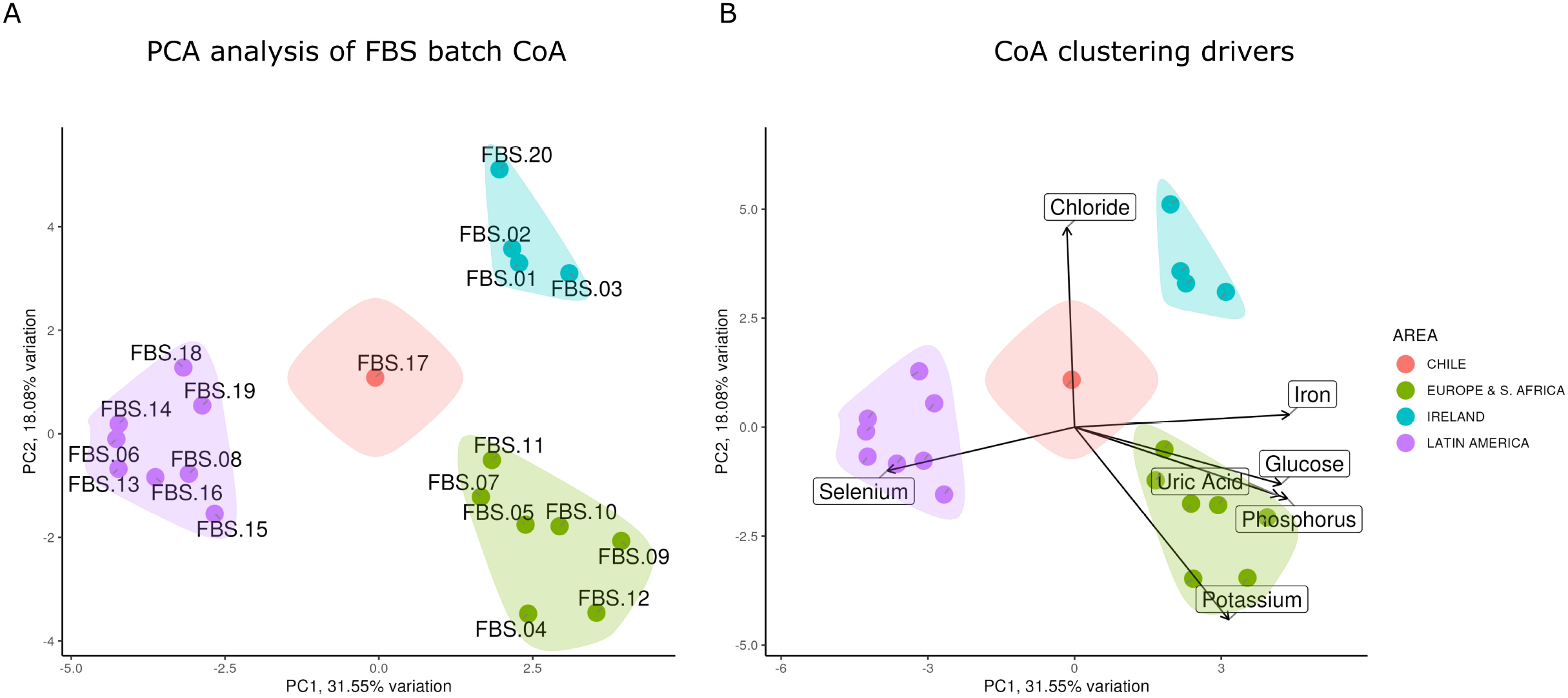
FBS biochemical profiles cluster by geographical origin. (A) Sample clustering by origin. PCA analysis was performed on chemical analysis of each FBS batch. The two first component are plotted (biplot) and colored by geographical origin with confidence ellipses. (B) Component loadings. PCA biplot showing variable loadings (arrows) overlaid on sample distribution. Loading vectors indicate the contribution of each biochemical parameter to the principal components, revealing which factors drive geographical patterns.

The correlation between CoAs and the transcriptome profiles based on an ssGSEA analysis, a useful method to compare samples across conditions, revealed that identical pathways exhibited different correlations with the same biochemical parameters. Nevertheless, in MRC-5 cells, few correlations were observed. Notably, selenium, inversely correlated with both pro-inflammatory cytokines and hallmarks of inflammatory responses, while uric acid was positively correlated. Coherently with the observed correlation between cytokines and CoAs, few correlations between transcriptome profiles and CoA parameters important for origin clustering were observed for Jurkat and THP-1 cells (Figure S3B). We conclude that taking all cell lines together, the CoA parameters do not correlate consistently with cytokine levels and transcriptome profiles and therefore, since the CoA determines the clustering of serum accordingly to origin, FBS batch origin seems not to constitute a good classifier for a general batch performance in relation to secretion of cytokines or transcriptome patterns.

## Discussion

FBS batch variability remains a persistent challenge for the scientific community, affecting both experimental reproducibility and cost efficiency. Traditionally, FBS batch selection relies primarily on the origin of production, a parameter widely used to classify FBS in terms of quality and price. Subsequent in-laboratory screening of multiple batches is then required to identify one that is compatible with previous experiments. In this study, we demonstrate that CoA parameters allow clear stratification of FBS according to origin; however, we also show that origin alone is not a universally informative predictor of batch performance. Although some CoA parameters correlated with biological outcomes - such as specific transcriptomic profiles and levels of secreted cytokines - these associations were not consistently observed across the different cell lines examined. This underscores the need for more comprehensive criteria than those currently included in CoAs for meaningful FBS classification. Additional parameters, including species background, vaccination history, feed composition, fetal age, and processing methods, may better account for the variation in FBS batch performance. Nevertheless, molecular-level approaches appear particularly promising for refining FBS characterization. One such approach is transcriptome profiling, which we propose as a robust and reproducible method capable of providing deeper insight into batch-specific effects.

Using three commonly employed cell lines - MRC-5, Jurkat, and THP-1 - and 20 FBS batches from different origins, we show that distinct batches induce cell line-specific patterns of differentially expressed genes. Gene set enrichment analysis further revealed that multiple pathways are differentially affected by FBS batches, with inflammatory and immune-related pathways showing particularly pronounced variability. These observations were corroborated by analyses of cytokines secreted by the three cell lines in response to the different FBS batches. Weighted Gene Co-expression Network Analysis identified co-expression modules that enabled additional resolution for comparing and stratifying FBS batches, including the identification of cell type–specific module clusters reflecting distinct molecular programs influenced by FBS batch differences.

Collectively, our findings show that the commonly held assumption that FBS origin is a reliable indicator of quality and performance does not hold when more comprehensive molecular analyses are applied. Moreover, our results support transcriptome profiling as a powerful tool for evaluating the performance of FBS batches and suggest its potential utility for selecting new batches to replace previously used ones.

This study is subject to limitations that may affect the interpretation and generalizability of the findings. First, the study was conducted using a limited number of cell lines, which are generally considered relatively robust to variations in FBS batches. Thus, inclusion of cell lines known to be more susceptible to FBS composition might reveal much stronger and cell line-dependent responses to different FBS batches. Consequently, testing a broader panel of cell lines derived from different tissues and even cultures of primary cells will be important for fully assessing the potential of transcriptome profiling to classify different types of FBS and to characterize batch-specific effects. Second, the number of serum batches analyzed in this study was limited. Evaluating a larger set of FBS samples from additional providers and of different origins would strengthen conclusions regarding the robustness of transcriptome classification and cytokines profiling in response FBS variability.

## Conclusion

In conclusion, our findings demonstrate that transcriptome profiling of cells cultured in different FBS batches of different origins represents a novel and sensitive approach for evaluating FBS performance. Notably, we show that geographic origin is not a reliable discriminator of FBS quality or functional impact across cell lines. The implementation of standardized molecular methods for FBS classification could greatly improve the selection of serum for experiments in which long-term reproducibility is critical, including applications in medical testing. Moreover, molecular profiling may ultimately facilitate the identification of suitable replacement batches when previously used FBS supplies are depleted, thereby reducing the time and costs associated with batch-to-batch variability in research.

## Data and materials availability

RNA-seq data have been deposited in the NCBI Gene Expression Omnibus (GEO) under accession number GSE314003. The data, currently in private mode, can be reviewed at the link https://www.ncbi.nlm.nih.gov/geo/ with the accession number and the token cvuzgccavfmhhgt. Analysis scripts to recreate the figures are available at https://gitlab.com/adlin-science-public/fbs-molecular-classification.git

## Acknowledgements

Adipocyte differentiation images were collected at the Center for Advanced Bioimaging (CAB) Denmark, University of Copenhagen.

## Funding declaration

This study was partially financed by Danmarks Innovationsfond (Innobooster), grant number 2055-00766A.

## Author contribution

Conceptualization: L.M., T.K., and K.K.; writing original draft: L.M., N.P.C., E.L., and K.K.; wet lab experiments: L.M., N.P.C., L.B., S.T.; data analysis and visualization: L.M., E.L., Q.S.; resources and supervision: T.K. and K.K.; all authors participated in discussions and contributed to the revision of the manuscript. All authors read and approved the final manuscript.

## Declaration of interest

L.M., N.P.C., T.K. are employees of Sera Scandia. S.T. is employee of BIOWEST. E.L. is employee of ADLIN Science. The other authors declare no competing interests.

## Supplemental information

Supplementary Information. Figures S1-S3, Tables S1.

Supplementary Data 1. Table S2. Differential expressed gene summary across FBS pairwise comparisons. Related to Figure 2A.

Supplementary Data 2. Table S3. Hallmark pathway enrichment by FBS batch comparison. Related to Figure 2B.

