## Supplementary information for "Towards molecular-based functional classification of fetal bovine serum"

Figure S1

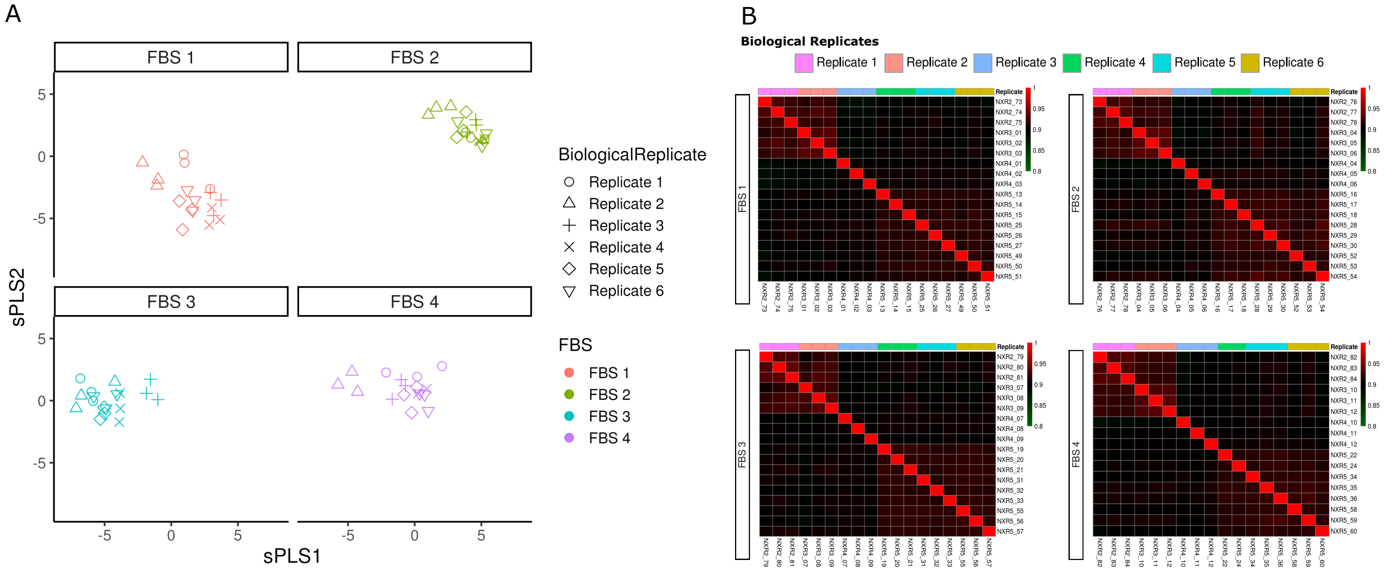


**Assessment of reproducibility in the FBS screening workflow.**

(A) Sparse Partial Least Squares (sPLS) analysis. SPLS plot of the first two components for samples grouped by FBS batches. Analysis is performed on VST-normalized data.
(B) Sample Correlation heatmap. Pearson correlation matrices within each FBS batch are calculated from log2-transformed TPM data.

Figure S2

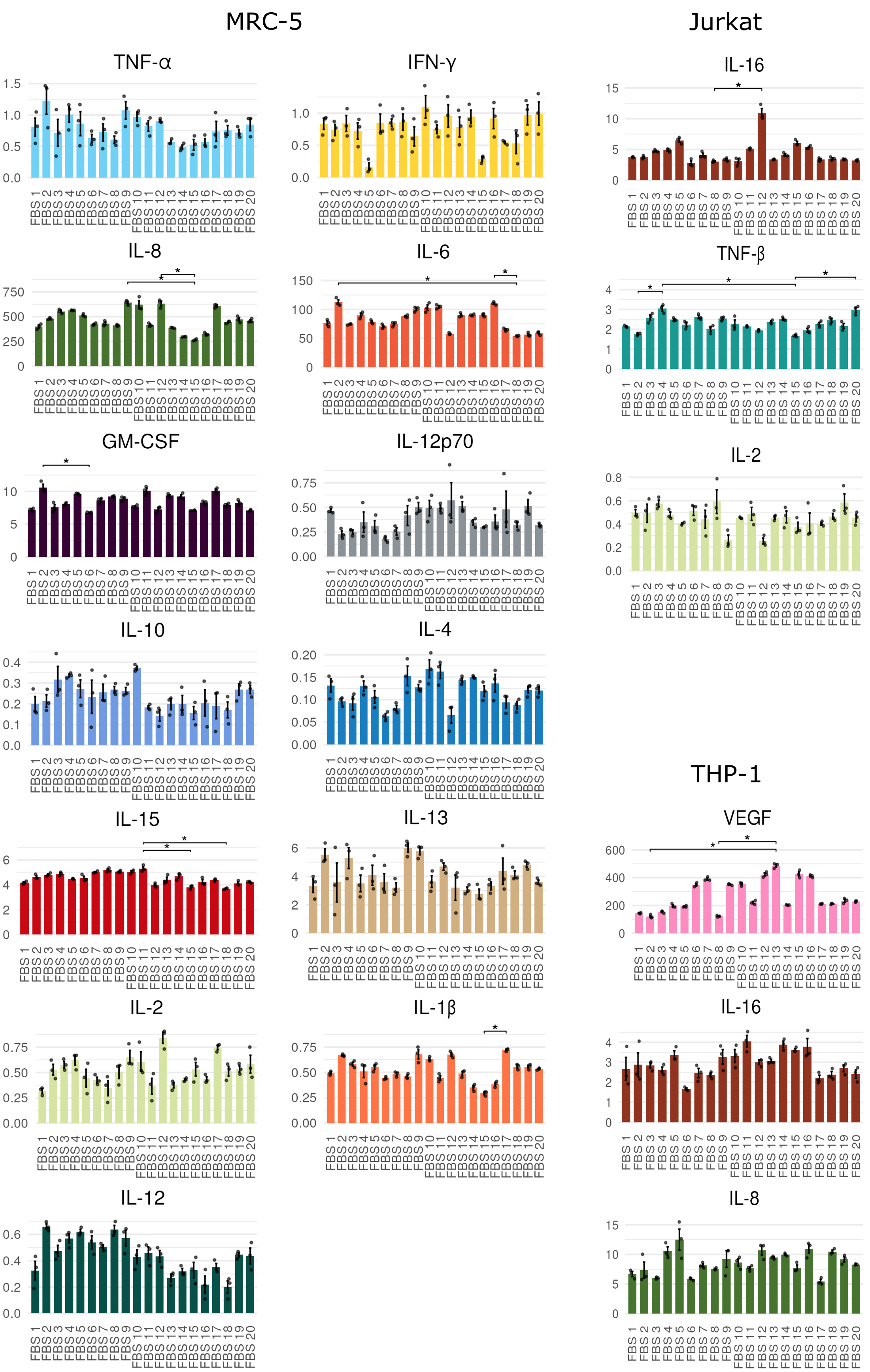


**Cytokine secretion profiles across FBS batches.**

FBS batch affects cytokine secretion in cell type-specific manner. Cytokine abundance measured in culture supernatants for three cell types cultured with different FBS batches. Bar plots show mean ± SEM for each cytokine. Individual points represent technical replicates. Statistical comparisons performed using Dunn's test with FDR correction. Significant differences (FDR < 0.05) are indicated by *

Figure S3

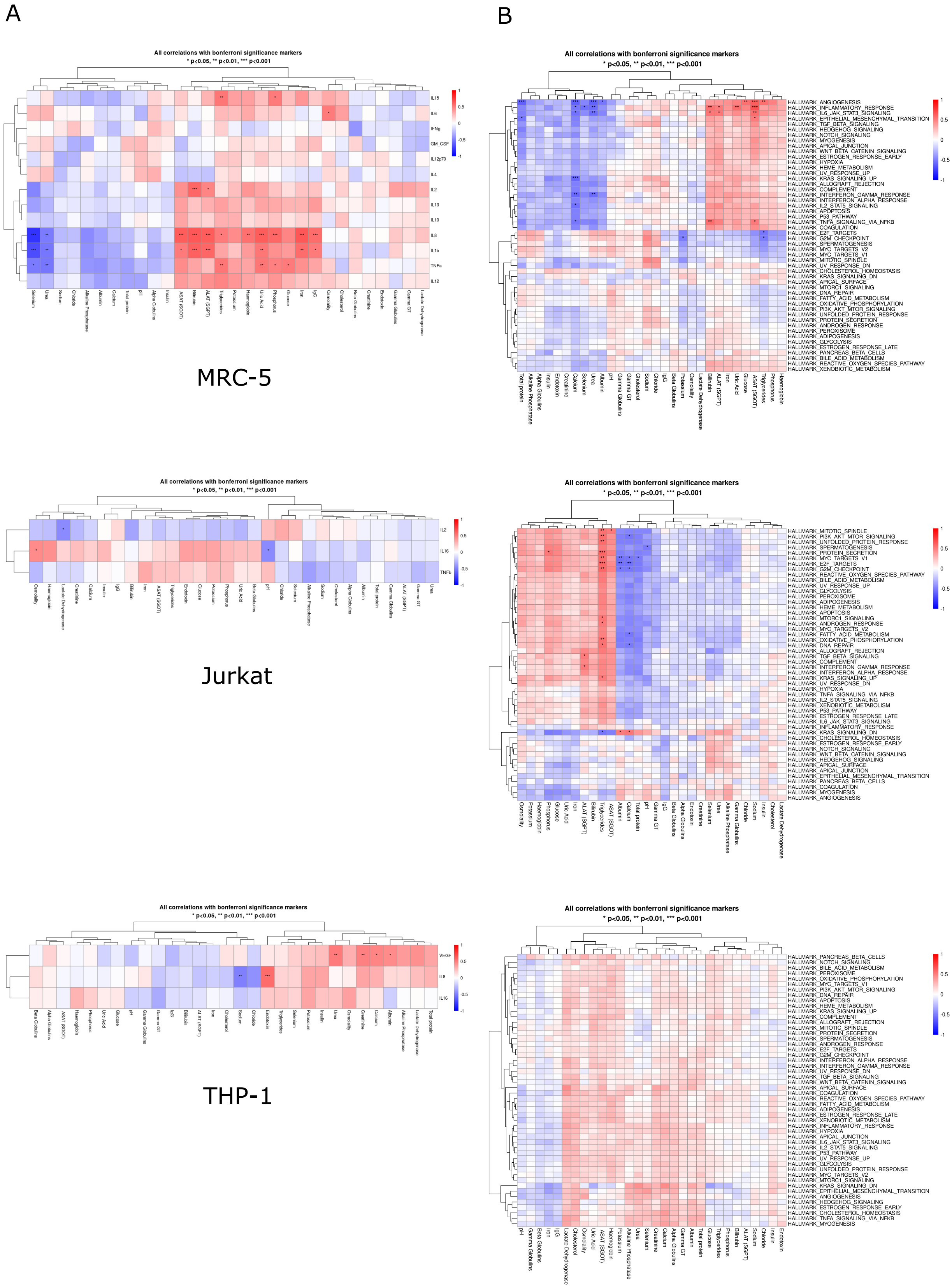


**Certain FBS biochemical parameters correlate with cytokine levels and hallmark pathway activities in a cell type-specific manner.**

(A) Correlation heatmap showing Spearman correlation coefficients between cytokine secretion levels and FBS biochemical composition parameters for each cell type.
(B) Correlation heatmaps showing Spearman correlations between GSVA pathway enrichment scores and FBS biochemical parameters for each cell type. GSVA scores are calculated from log2 (TPM + 0.001) expression data using MSigDB Hallmark gene sets.
Color scale indicates in blue negative correlation and in red positive correlation. Significative correlations after Bonferroni correction are indicated as *p < 0.05, **p < 0.01, ***p < 0.001.

Table S1. **Parameters of FBS batches.**

| Info | Units | FBS 1 | FBS 2 | FBS 3 | FBS 4 | FBS 5 | FBS 6 | FBS 7 | FBS 8 | FBS 9 | FBS 10 |
|---|---|---|---|---|---|---|---|---|---|---|---|
| **BATCH** | | S00MZ2 | S00JI | S00E11 | S00E9101 | S00E71 | S00CM1 | S00GF1 | S00L81 | S00MF1 | S00LN2 |
| **ORIGIN** | | IRELAND | IRELAND | IRELAND | SPAIN | SPAIN | PANAMA | S. AFRICA | PARAGUAY | FRANCE | FRANCE |
| **VALIDATION DATE** | | 13/09/2022 | 30/09/2021 | 30/01/2020 | 13/02/2020 | 13/02/2020 | 12/09/2019 | 03/11/2020 | 24/02/2022 | 07/07/2022 | 07/04/2022 |
| **EXPIRY DATE** | | 12/09/2027 | 29/09/2026 | 28/01/2025 | 11/02/2025 | 11/02/2025 | 10/09/2024 | 02/11/2025 | 23/02/2027 | 06/07/2027 | 06/04/2027 |
| **pH** | | 7,36 | 7,28 | 7,42 | 7,36 | 7,33 | 7,29 | 7,30 | 7,56 | 7,34 | 7,32 |
| **Osmolality** | mOsm/kg | 309 | 310 | 304 | 315 | 304 | 305 | 316 | 308 | 314 | 313 |
| **Endotoxin** | EU/ml | 0,376 | < 0,200 | 0,257 | 16.370 | 2.809 | 0,398 | 1.035 | 1.221 | 0,640 | 0,557 |
| **Haemoglobin** | mg/100ml | 12,75 | 15,14 | 25,92 | 19,29 | 18,92 | 11,89 | 12,43 | 14,64 | 23,67 | 18,41 |
| **Total protein** | g/l | 33,6 | 33,1 | 32,3 | 37,2 | 35,9 | 35,5 | 37,7 | 36,3 | 35,4 | 35,7 |
| **ALAT (SGPT)** | IU/l | 6 | 8 | 11 | 6 | 7 | 8 | <6 | 9 | 9 | 9 |
| **Alkaline Phosphatase** | IU/l | 233 | 210 | 190 | 228 | 202 | 346 | 229 | 329 | 253 | 253 |
| **ASAT (SGOT)** | IU/l | 33 | 45 | 59 | 51 | 42 | 27 | 46 | 26 | 67 | 58 |
| **Bilirubin** | mg/100ml | 0,18 | 0,21 | 0,22 | 0,19 | 0,22 | 0,16 | 0,16 | 0,21 | 0,25 | 0,21 |
| **Calcium** | mg/100ml | 12,4 | 12,5 | 12,7 | 14,0 | 13,9 | 14,2 | 13,4 | 13,3 | 13,4 | 13,4 |
| **Gamma GT** | IU/l | 6 | 6 | 6 | 7 | 6 | 7 | 5 | 7 | 6 | 6 |
| **Cholesterol** | mg/100ml | 30,0 | 32,0 | 33,0 | 33,0 | 30,0 | 34,0 | 31,0 | 35,0 | 33,0 | 33,0 |
| **Creatinine** | mg/100ml | 2,5 | 2,3 | 2,2 | 2,9 | 2,7 | 2,8 | 2,7 | 2,7 | 2,8 | 2,7 |
| **Chloride** | mmol/l | 101 | 105 | 103 | 99 | 96 | 103 | 98 | 98 | 98 | 98 |
| **Glucose** | mg/100ml | 151,0 | 133,0 | 104,0 | 147,0 | 128,0 | 60,0 | 167,0 | 69,0 | 137,0 | 129,0 |
| **Iron** | µg/100ml | 201 | 189 | 208 | 199 | 195 | 153 | 215 | 156 | 209 | 204 |
| **Lactate Dehydrogenase** | IU/l | 393 | 366 | 341 | 605 | 541 | 519 | 506 | 518 | 621 | 582 |
| **Phosphorus** | mg/100ml | 10,0 | 10,1 | 10,2 | 10,5 | 10,3 | 9,0 | 11,4 | 9,0 | 11,3 | 11,1 |
| **Potassium** | mmol/l | 11,3 | 11,8 | 11,9 | 16,1 | 15,1 | 11,3 | 13,7 | 11,7 | 14,9 | 14,7 |
| **Sodium** | mmol/l | 136 | 138 | 134 | 133 | 128 | 137 | 136 | 132 | 133 | 133 |
| **Triglycerides** | mg/100ml | 67,0 | 69,0 | 69,0 | 77,0 | 75,0 | 62,0 | 78,0 | 66,0 | 88,0 | 89,0 |
| **Urea** | mg/100ml | 22,0 | 26,0 | 28,0 | 31,0 | 30,0 | 40,0 | 35,0 | 40,0 | 34,0 | 33,0 |
| **Uric Acid** | mg/100ml | 3,2 | 2,8 | 2,2 | 3,3 | 2,9 | 1,5 | 2,6 | 1,9 | 3,2 | 3,0 |
| **Albumin** | g/l | 14,5 | 14,1 | 13,6 | 15,7 | 15,4 | 17,3 | 16,9 | 15,9 | 15,5 | 15,3 |
| **Alpha Globulins** | g/l | 13,2 | 13,5 | 12,8 | 15,5 | 13,1 | 16,4 | 13,3 | 18,2 | 12,8 | 15,3 |
| **Beta Globulins** | g/l | 5,5 | 5,1 | 5,5 | 5,4 | 7,1 | 1,8 | 6,9 | 2,0 | 6,7 | 4,6 |
| **Gamma Globulins** | g/l | 0,4 | 0,4 | 0,5 | 0,5 | 0,4 | 0,1 | 0,5 | 0,6 | 0,4 | 0,5 |
| **IgG** | mg/l | 107,0 | 176,0 | 159,1 | 186,9 | 131,3 | 119,1 | 114,3 | 86,0 | 83,0 | 377,0 |
| **Gamma Irradiated** | Yes/No | No | No | No | No | No | No | No | No | No | No |
| **Selenium** | µg/l | 16 | 22 | 18 | 19 | 18 | 22 | 26 | 31 | 15 | 17 |
| **Insulin** | µUI/mL | 61,5 | 90,6 | 115 | 107 | 148 | 147 | 112 | 113 | 101 | 110 |

| Info | Units | FBS 11 | FBS 12 | FBS 13 | FBS 14 | FBS 15 | FBS 16 | FBS 17 | FBS 18 | FBS 19 | FBS 20 |
|---|---|---|---|---|---|---|---|---|---|---|---|
| **BATCH** | | S001U2 S13992S1820 | S00132 S13119S1820 | S00GT1 | S00ML1 | S0017201 S13205S181G | S00L4201 | S00OJ | S00PB | S00P0 | S00OZ |
| **ORIGIN** | | FRANCE | FRANCE | COLOMBIA | COLOMBIA | COLOMBIA | COLOMBIA | CHILE | GUATEMALA | GUATEMALA | IRELAND |
| **VALIDATION DATE** | | 06/12/2021 | 29/05/2020 | 10/12/2020 | 09/08/2022 | 06/12/2021 | 22/02/2022 | 03/02/2023 | 20/04/2023 | 06/04/2023 | 04/04/2023 |
| **EXPIRY DATE** | | 05/11/2023 | 19/03/2023 | 09/12/2025 | 08/08/2027 | 14/04/2023 | 21/02/2027 | 02/02/2028 | 18/04/2028 | 04/04/2028 | 02/04/2028 |
| **pH** | | 7,36 | 7,05 | 7,54 | 7,46 | 7,14 | 7,45 | 7,98 | 7,69 | 7,80 | 7,49 |
| **Osmolality** | mOsm/kg | 320 | 330 | 308 | 307 | 316 | 308 | 299 | 292 | 302 | 299 |
| **Endotoxin** | EU/ml | 1.313 | 7.225 | 0,406 | 1.383 | 1.196 | 0,385 | 0,302 | 1.566 | 0,853 | 0,488 |
| **Haemoglobin** | mg/100ml | 21,94 | 24,77 | 12,86 | 10,90 | 15,29 | 14,33 | 15,56 | 4,89 | 11,30 | 11,70 |
| **Total protein** | g/l | 35,2 | 34,9 | 37,5 | 36,9 | 36,5 | 38,1 | 38,0 | 37,4 | 38,1 | 31,6 |
| **ALAT (SGPT)** | IU/l | <6 | 12 | <6 | <6 | <6 | 6 | 8 | <6 | <6 | 6 |
| **Alkaline Phosphatase** | IU/l | 168 | 241 | 253 | 273 | 361 | 334 | 225 | 290 | 298 | 195 |
| **ASAT (SGOT)** | IU/l | 11 | 46 | 20 | 12 | 35 | 29 | 32 | 17 | 17 | 44 |
| **Bilirubin** | mg/100ml | 0,1 | 0,22 | 0,13 | 0,10 | 0,19 | 0,18 | 0,23 | 0,16 | 0,10 | 0,23 |
| **Calcium** | mg/100ml | 13,5 | 13,9 | 14,1 | 14,0 | 14,1 | 13,8 | 13,8 | 14,0 | 13,8 | 12,4 |
| **Gamma GT** | IU/l | 5 | 6 | 5 | 5 | 7 | 7 | 9 | 6 | 6 | 6 |
| **Cholesterol** | mg/100ml | 30,0 | 31 | 32,0 | 32,0 | 33 | 35,0 | 33,0 | 29,0 | 32,0 | 30,0 |
| **Creatinine** | mg/100ml | 2,7 | 2,9 | 2,7 | 2,7 | 2,9 | 2,7 | 2,6 | 2,7 | 2,8 | 2,0 |
| **Chloride** | mmol/l | 98 | 98 | 99 | 100 | 100 | 100 | 99 | 97 | 100 | 103 |
| **Glucose** | mg/100ml | 129 | 140 | 86,0 | 74,0 | 90 | 73,0 | 99,0 | 71,0 | 88,0 | 94,0 |
| **Iron** | µg/100ml | 211 | 202 | 157 | 166 | 147 | 151 | 213 | 171 | 175 | 191 |
| **Lactate Dehydrogenase** | IU/l | 338 | 585 | 388 | 314 | 687 | 598 | 510 | 439 | 431 | 367 |
| **Phosphorus** | mg/100ml | 10,7 | 10,5 | 9,1 | 9,2 | 9,5 | 8,8 | 10,1 | 8,5 | 9,2 | 9,9 |
| **Potassium** | mmol/l | 13,7 | 14,1 | 12,0 | 11,7 | 12,0 | 11,5 | 11,4 | 11,1 | 12,2 | 11,9 |
| **Sodium** | mmol/l | 132 | 133 | 134 | 135 | 136 | 135 | 136 | 131 | 135 | 134 |
| **Triglycerides** | mg/100ml | 82,0 | 84 | 67,0 | 60,0 | 59 | 62,0 | 52,0 | 55,0 | 64,0 | 70,0 |
| **Urea** | mg/100ml | 35,0 | 32 | 43,0 | 42,0 | 41 | 40,0 | 35,0 | 31,0 | 33,0 | 23,0 |
| **Uric Acid** | mg/100ml | 2,6 | 3,0 | 1,8 | 1,7 | 2,0 | 1,9 | 3,1 | 1,6 | 1,9 | 2,0 |
| **Albumin** | g/l | 14,2 | 14,5 | 17,3 | 15,8 | 16,6 | 16,9 | 16,6 | 18,2 | 16,5 | 13,2 |
| **Alpha Globulins** | g/l | 14,8 | 15,4 | 18,1 | 19,0 | 13,6 | 15,2 | 15,2 | 11,0 | 15,1 | 13,1 |
| **Beta Globulins** | g/l | 5,9 | 4,4 | 1,9 | 1,8 | 5,8 | 5,5 | 5,6 | 7,6 | 6,0 | 4,9 |
| **Gamma Globulins** | g/l | 0,3 | 0,5 | 0,2 | 0,3 | 0,5 | 0,6 | 0,6 | 0,6 | 0,5 | 0,4 |
| **IgG** | mg/l | 222,0 | 119,7 | 90,0 | 72,0 | 12,7 | 101,0 | 568,0 | 134,0 | 152,0 | 53,0 |
| **Gamma Irradiated** | Yes/No | No | No | No | No | Yes | Yes | No | No | No | No |
| **Selenium** | µg/l | 20 | 14 | 37 | 29 | 33 | 31 | 13 | 23 | 23 | 16 |
| **Insulin** | µUI/mL | 127 | 159 | 139 | 141 | 84,5 | 125 | 64,5 | 111 | 115 | 94,2 |
